# Comparative analysis of the RNA-chromatin interactions data. Completeness and accuracy

**DOI:** 10.1101/2023.09.21.558854

**Authors:** Grigory Ryabykh, Artem Vasilyev, Lidia Garkul, Vladimir Shatskiy, Andrey Mironov

## Abstract

Non-coding RNAs play an essential role in a wide variety of biological processes. We have previously developed RNA-Chrom, an analytical database containing uniformly processed RNA-chromatin interactions data. Here, we analyzed the consistency of these data with each other and performed a comparative analysis of human and mouse datasets from “all-to-all” experiments (interactome of all possible RNAs in a cell) and “one-to-all” experiments (individual RNA interactome). We analyzed the dependence of RNA cis-contacts density as a function of distance to the RNA source gene and found that in all experiments, density of cis-contacts decreases with distance. We tested whether the most contacting RNAs of one “all-to-all” experiment (“RNA-leaders”) are the same for the other “all-to-all” experiments. Our analysis shows that “all-to-all” experiments are not complete and require substantially deeper sequencing, and conclusions can only be drawn for highly contacting RNAs. We noted that for “all-to-all” experiments, sequencing depth and the type of cell treatment give a significant impact on the contact map, while the cell line does not have as much of an impact.

## 1. Introduction

Increasing amounts of evidence is emerging every year that non-coding RNAs (ncRNAs) in animals and plants are involved in a wide variety of biological processes such as regulation of cell differentiation, development, gene expression, chromatin remodeling, and maintenance of chromatin structure, splicing, RNA processing, formation of biomolecular condensates, and others. In addition, disruption of regulatory pathways mediated by ncRNAs may lead to various diseases [1].

RNA molecules interact with a variety of proteins, chromatin and other RNAs. Experimental methods have been developed to identify DNA loci contacted by ncRNAs, which can be categorized into two groups: «one-to-all» (or OTA) and «all-to-all» (or ATA). The first group of methods (RAP [2], CHART-seq [3], ChIRP-seq [4], dChIRP-seq [5], ChOP-seq [6], CHIRT-seq [7]) determines the contacts of a previously known RNA with chromatin, while the second group of methods (MARGI [8], GRID-seq [9], ChAR-seq [10], iMARGI [11], RADICL-seq [12], Red-C [13]) aims to determine all possible RNA-DNA contacts in a cell [14].

The RNA-Chrom database [15] has collected a large amount of uniformly processed RNA-chromatin interactions data. The variety of cell lines on the one hand forms a diverse picture of the behavior of ncRNAs in human and mouse cells. On the other hand, for almost every ncRNA, researchers chose a unique cell line and specific experimental conditions (Suppl. Tables 1 and 2), which makes the picture fragmented and their comparison difficult. In the present study, we performed a comparative analysis of the RNA-chromatin interactions data from the RNA-Chrom database. We assessed data consistency, accuracy, the level of noise in it, the effect of the biological material preparation. We analyzed the dependence of RNA cis-contacts density on the distance between the gene from which this RNA is transcribed (hereafter referred to as “RNA source gene”) and chromatin target loci. By analogy with the chromatin-chromatin interactome data, we will refer to this dependence as “scaling”. Moreover, an interspecies comparison of the data was performed.

## 2. Materials and Methods

### 2.1. Data

RNA-chromatin interactions data, excluding “singletons”, were taken from RNA-Chrom database [15]: without normalization to background (“raw”), normalized to background (“normalized”), and background-normalized contacts crossing MACS2 peaks. All experimental identification numbers (“Exp.IDs”) referenced in this paper correspond to the identifiers from the RNA-Chrom DB (Suppl. Tables 3 and 4). OTA experiments were accompanied by an input that was taken as a background. Since no additional data with background (non-specific) contacts are provided for ATA data, we used an endogenous background model on mRNA trans-contacts [9]. In cases when RNA source genes were required for analysis, we excluded experiments on multicopy RNAs (*Homo sapiens*: 7SK, ERV-9 RNAs; *Mus musculus*: TERRA, PAR-TERRA, 7SK, U1, LINE1, IAPEz-int, IAP, 116HG, B2 RNAs).

### 2.2. Correlation analysis

We calculated the total contacts tracks of all RNAs for raw and normalized data. To do this, each chromosome was divided into bins of size 10 kb and 100 kb. For each dataset for individual RNA, all the contacts across these bins were counted. Thus, for each experiment, replicates of the experiment, or individual RNA, we obtained the contact counts vector across the bins. Then we calculated pairwise Pearson correlations between these vectors. The contacts vector for an experiment is the sum of the contacts vectors of all its replicates. No multicopy RNA tracks were used in calculating correlations between OTA and ATA experiments except for 7SK and U1, because for the remaining multicopy RNAs, genes in ATA were not identified.

To ensure that the RNA contacts overrepresentation near the RNA source gene does not affect the results, all analyses excluded contacts within 100 kb of the RNA source gene, except in cases where the RNA originated from multicopy genes.

### 2.3. Chromatin potential

In papers dealing with ATA experiments [9,12,13] it was noted that the higher the RNA expression is, the more contacts are observed. To distinguish RNAs that are more prone to form contacts with chromatin, we introduced the concept of “chromatin potential”. For each RNA, this value is calculated with the following equation:

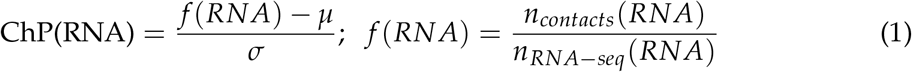

where *μ* is the average *f* (*RNA*) for all RNAs, *σ* is the standard deviation. The chromatin potential value characterizes the excess of the number of observed contacts of a given RNA over the expected number of contacts. To calculate the chromatin potential, total ribo-minus RNA-seq data were taken: GSM4041596 and GSM4041597 for the human K562 cell line, GSM2400249 and GSM2400250 for the mouse embryonic stem cell line.

### 2.4. Scaling

Several studies on ATA [12,13] noted that the RNA cis-contacts density decreases with distance from the RNA source gene. Unlike the correlation analysis, in scaling analysis we binned distances according to the geometric law:

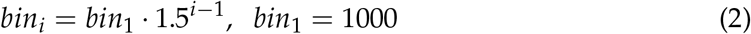

To ensure comparability of results from different experiments, we normalized the density by the total number of cis-contacts of the corresponding track:

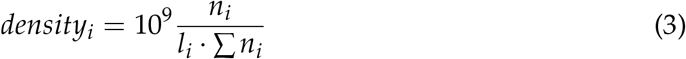

where *n*_*i*_ is the number of background-normalized cis-contacts of the given RNA or the considered set of RNAs in the *i*-th bin, and *l*_*i*_ is the *i*-th bin length.

In the case of ATA data, we considered the 25% of RNAs with the highest number of cis-contacts (Suppl. Table 5) to avoid the influence of RNAs with a small number of cis-contacts.

We divided mouse and human OTA experiments into three groups. For mouse: Xist RNA (47 experiments), Malat1 RNA (12 experiments) and all other mouse RNAs (28 experiments, 20 RNAs). For human: MALAT1 RNA (10 experiments), NEAT1 RNA (10 experiments) and all other human RNAs (25 experiments, 20 RNAs).

### 2.5. Search for orthologs

RNA orthologs were established using the ortho2align program [16]. The orthologs obtained using this algorithm were further selected based on length – we filtered out too short orthologs (length less than 40% of the query).

## 3. Results

### 3.1. Analysis of replicates

To ensure the validity of OTA and ATA data, each experiment was performed several times, so that several independent replicates were obtained for the same conditions. If there is a high correlation between replicates of one experiment, the results of the experiment can be considered reliable. Lack of correlation between replicates indicates a high level of noise in the data, as well as insufficient sequencing depth, due to which poorly overlapping of the true signal in the replicates can be seen, which together does not allow us to interpret the results of the experiment unambiguously and may indicate that the data are incomplete. To test for replicate consistency, we calculated Pearson correlations between replicates (bin size: 10 kb) for those experiments from RNA-Chrom DB for which at least two replicates were made: 37 (of 53) OTA experiments for human and 40 (of 118) for mouse, 6 (of 9) ATA experiments for human and 7 (of 7) for mouse (Figure 1). In a large part of the experiments, the replicates are sufficiently well correlated (correlation >0.75), so further analysis of the corresponding data can be considered reliable. We noted that for all human ATA data, the replicate correlation is significantly lower. Therefore, we later used 100 kb bins when analyzing these data (Suppl. Figure 1).

**Figure 1.**
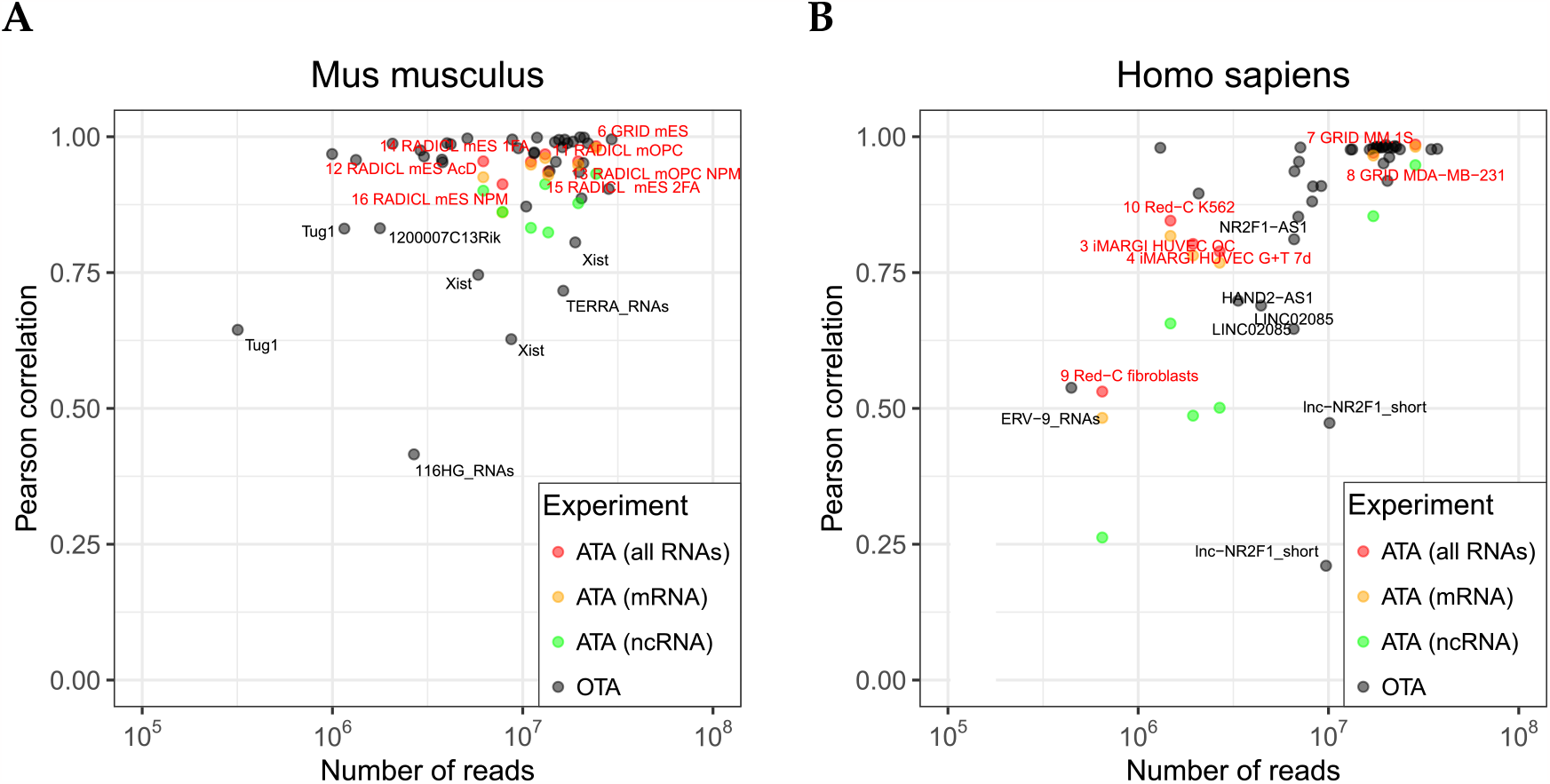
Pearson correlation between replicates as a function of the minimum number of raw contacts in them: (**A**) mouse OTA and ATA experiments; (**B**) human OTA and ATA experiments. If an experiment had more than two replicates, the average of all pairwise correlations was taken. Genomic bin size: 10 kb, contacts filtering: >100 kb from the RNA source gene.

For ATA experiments, the correlation of contacts in their replicates is higher for mRNAs than for ncRNAs. This is apparently due to the fact that the transcription level and total number of contacts of mRNAs are much higher than for ncRNAs (Figure 1). In the subsequent analysis, we merged the replicates.

In the analysis of ATA replicates, we compared total tracks of RNA-chromatin contacts as was done in [9,13] – the contacts of all RNAs were summed. Next, we compared contacts tracks in replicates of individual RNAs. To do this, we selected the top 5% mRNAs and top 5% ncRNAs in terms of the number of contacts in the “Red-C K562” (Exp.ID: 10, human) and “RADICL mES 1FA” (Exp.ID: 14, mouse) experiments (Figure 2). The figure indicates that when the number of contacts of an individual RNA in one of the replicates is less than 1000, the correlation is quite low (correlation <0.5). It can be also noticed that for mouse RNAs, a 100 kb bin greatly increases correlations, and for human RNAs, a 10 kb bin decreases correlations. So the reasoning of bin-size choice is to find a balance between accuracy and consistency. In this regard, in future analyses of mouse and human ATA data, we used 10 kb and 100 kb bins, respectively.

**Figure 2.**
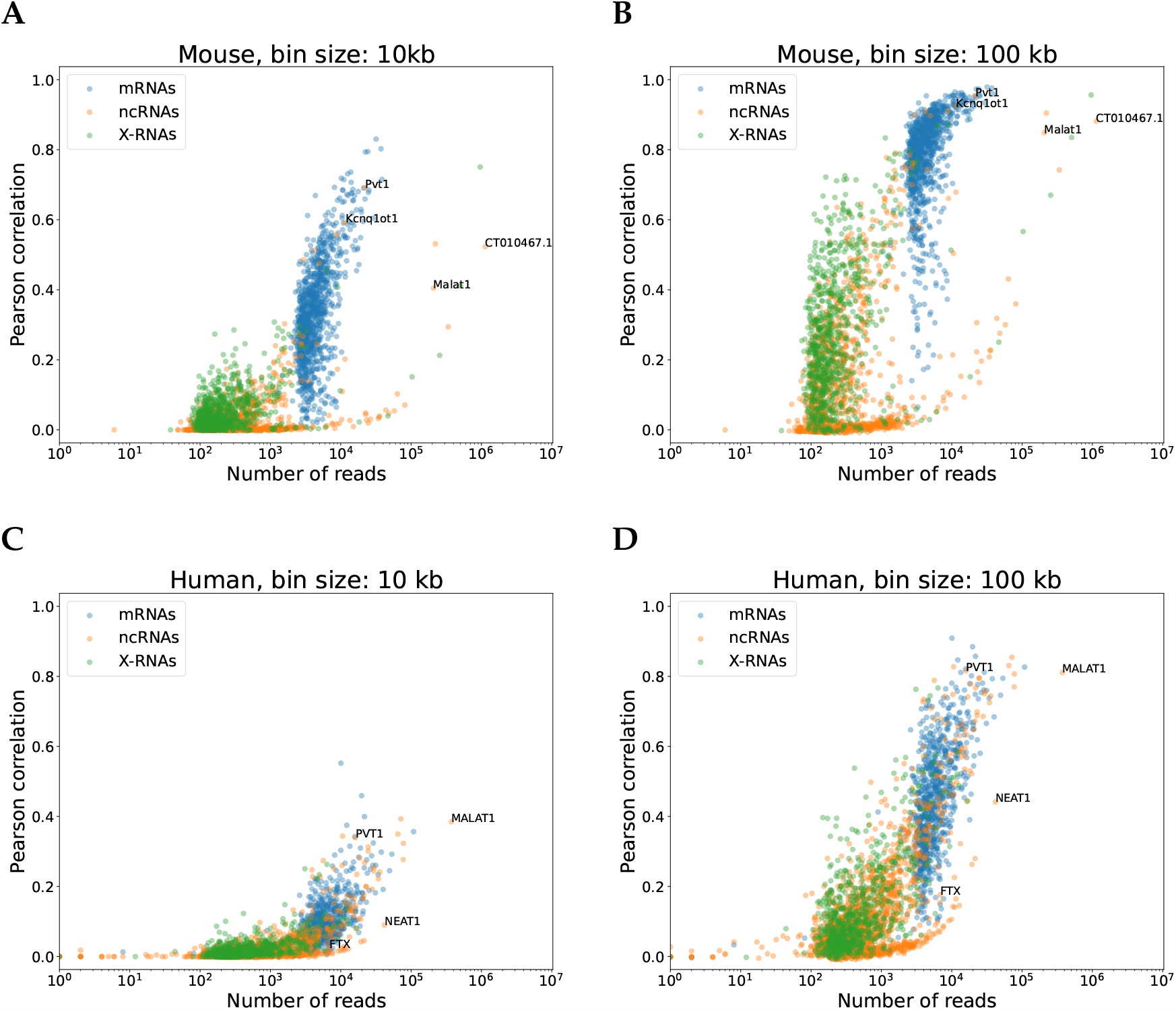
Pearson correlation between raw contacts tracks of RNAs in replicates as a function of the minimum number of contacts in them. Top 5% mRNAs and top 5% ncRNAs in terms of the number of contacts with chromatin. Contacts filtering: >100 kb from the RNA source gene. (**A, B**) Exp.ID: 14, “RADICL mES 1FA”, two largest replicates. (**C, D**) Exp.ID: 10, “Red-C K562”, two largest replicates. Genomic bin size: 10 kb and 100 kb.

### 3.2. Scaling

In the RNA-chromatin interactome data, the density of RNA contacts depends on the distance between the RNA source gene and the DNA target loci located on the same chromosome. Such contacts are further mentioned as “cis-contacts”, and this dependence as “scaling”. This dependence is similar to the scaling in the DNA-DNA interactome data (Hi-C methods) [17]. Scaling in the RNA-chromatin interactome is explained by the ratio of the rate of synthesis (and degradation) of RNA to its rate of diffusion through the nucleus, as well as by chromatin folding. With the minimum synthesis rate and rapid diffusion, this dependence is flat, and with the opposite ratio, the rate of density decline is large.

We compared the behavior of scaling in different experiments. In human and mouse ATA data, the scaling curves behave similarly and are independent of both the chromosome, from which the corresponding RNAs are transcribed, and the RNA biotype. The exception is the two mouse experiments that were treated with protease K (Exp.ID: 13, 16 – NPM-experiments; Figure 3). This behavior suggests that cis-contacts cover the chromatin more evenly in these data. The authors [12] suggest that a significant change in the number of contacts with distance is due to the fact that protease K treatment greatly reduced the share of the protein-mediated RNA-DNA contacts. The authors suggest that this treatment predominantly leaves only those contacts formed through structures such as RNA-DNA-DNA triplexes and R-loops. But the stability of heteroduplexes in R-loops in the absence of proteins seems questionable.

**Figure 3.**
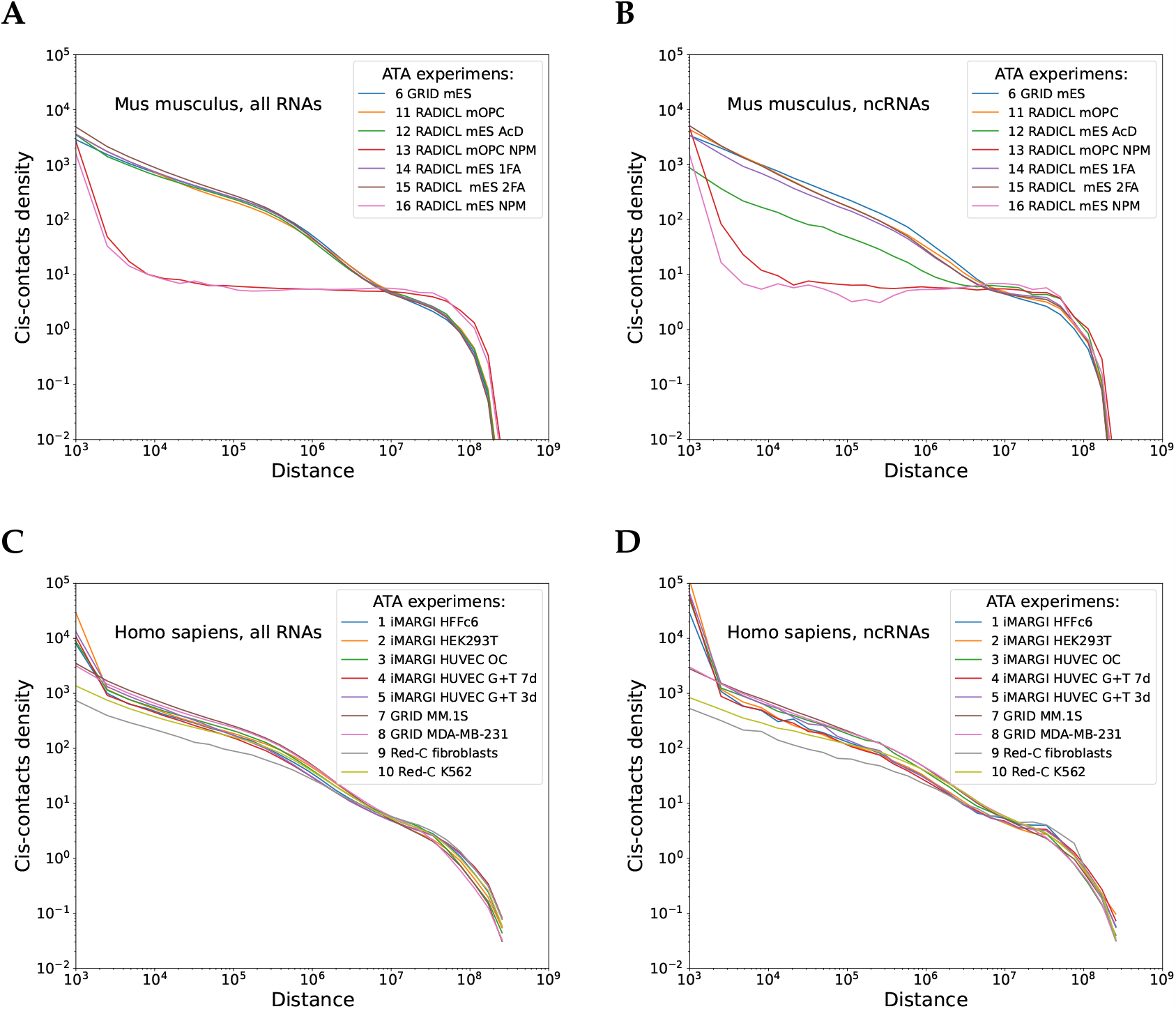
Scaling curves plotted for all (**A, B**) mouse and (**C, D**) human ATA data from the RNA-Chrom DB. Each curve is the scaling behavior for the corresponding dataset (see “Materials and Methods” section). The format of the ATA experiment labels: *“Exp.ID from RNA-Chrom _ cell line _ treatment”*. Treatment: “NPM” – proteinase K, “1FA” – 1% formaldehyde, “2FA” – 2% formaldehyde, “AcD” – actinomycin D, “OC” – osmotic control, “G+T 3d” – combining high glucose (to mimic hyperglycemia) and TNFalpha (to mimic inflammation) for 3 days, “G+T 7d” – combining high glucose (to mimic hyperglycemia) and TNFalpha (to mimic inflammation) for 7 days.

The scaling in OTA data looks slightly different if compared with ATA data (Figure 4). This can be explained as a consequence of the sequencing depth per RNA being much lower in ATA data as compared to OTA data. A variety of cell treatments also lead to changes in scaling.

**Figure 4.**
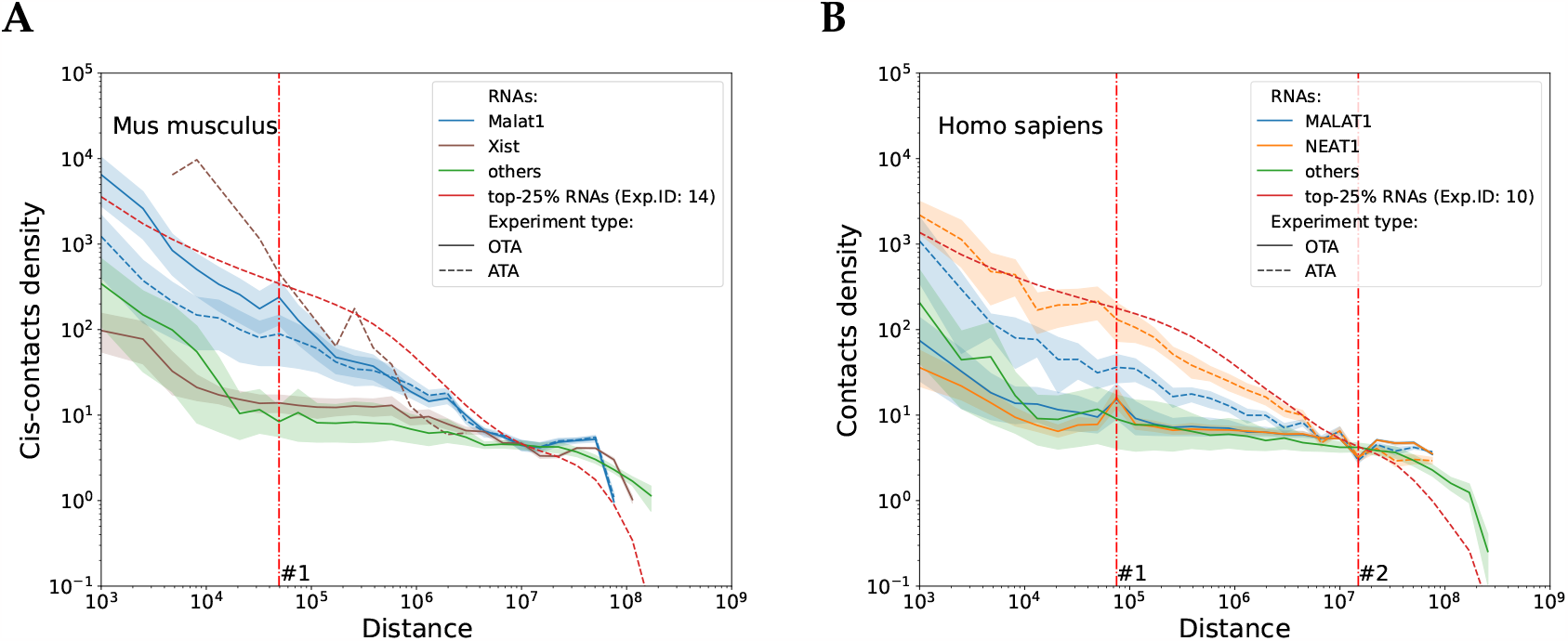
Average scaling curves plotted for all (**A**) mouse and (**B**) human OTA data. Different colors indicate the groups of RNAs considered for the analysis, solid line corresponds to OTA data, dashed line corresponds to ATA data. Red vertical lines: #1) peak of Malat1 RNA contacts density with Neat1 gene (mouse), and MALAT1 and NEAT1 RNAs with each other’s genes (human); #2) dip in contacts density for MALAT1 and NEAT1 RNAs at a distance that is equal to the distance to the centromeric region (human).

The MALAT1 and NEAT1 RNA scaling curves for OTA data, but not for ATA data, show a pronounced peak (Figure 4, vertical line #1). This peak reflects direct interaction of MALAT1 RNA with the NEAT1 gene and and vice versa: distance between these genes in mouse genome – 22037 bp, in human genome – 52149 bp. This observation corresponds to the results of the foregoing research [18]. These curves show also chromosomal feature for both OTA and ATA data: the human data show a dip at 11 Mb from the gene, which corresponds to the distance from RNA source gene to the centromeric region, (Figure 4, vertical line #2), while the mouse data do not have the dip, since the 19th chromosome is acrocentric.

In mouse and human OTA data, which we consider more reliable, the scaling curves for RNA from the “others” group reach a conditional plateau between 10 kb and 100 kb (Figure 4), which may suggest a “polymerase trace”. To summarize, scaling not only reflects the process of RNA diffusion from the transcription site, but also contains information about chromatin structure. In addition, an extra chemical treatment during data acquisition and sequencing depth per RNA influence the scaling.

### 3.3. Correlation analysis

#### 3.3.1. Comparison of OTA experiments

To investigate contacts consistency and possible common functions of different RNAs, and to estimate the overall noise, we compared RNA contacts tracks from different OTA experiments. To evaluate the role of noise, we applied the following filters: normalizing for background, cutting off proximal contacts (<100 kb from the RNA source gene) and accepting only those contacts that fall within the peaks defined by MACS2. Suppl. Figures 2A and 2B demonstrate how different filters affect the correlation. In raw data, the correlations between the contacts tracks of different RNAs are quite high. However, the application of filters improves the situation significantly. The correlations for different RNAs decrease slightly when normalized and significantly when moving to contacts from the peaks. Polymerase trace filtering does not affect the correlations in raw and normalized data, but markedly reduces the correlations for peaks. A more detailed picture of the RNA correlations is shown in Figure 5 and Suppl. Figure 3.

**Figure 5.**
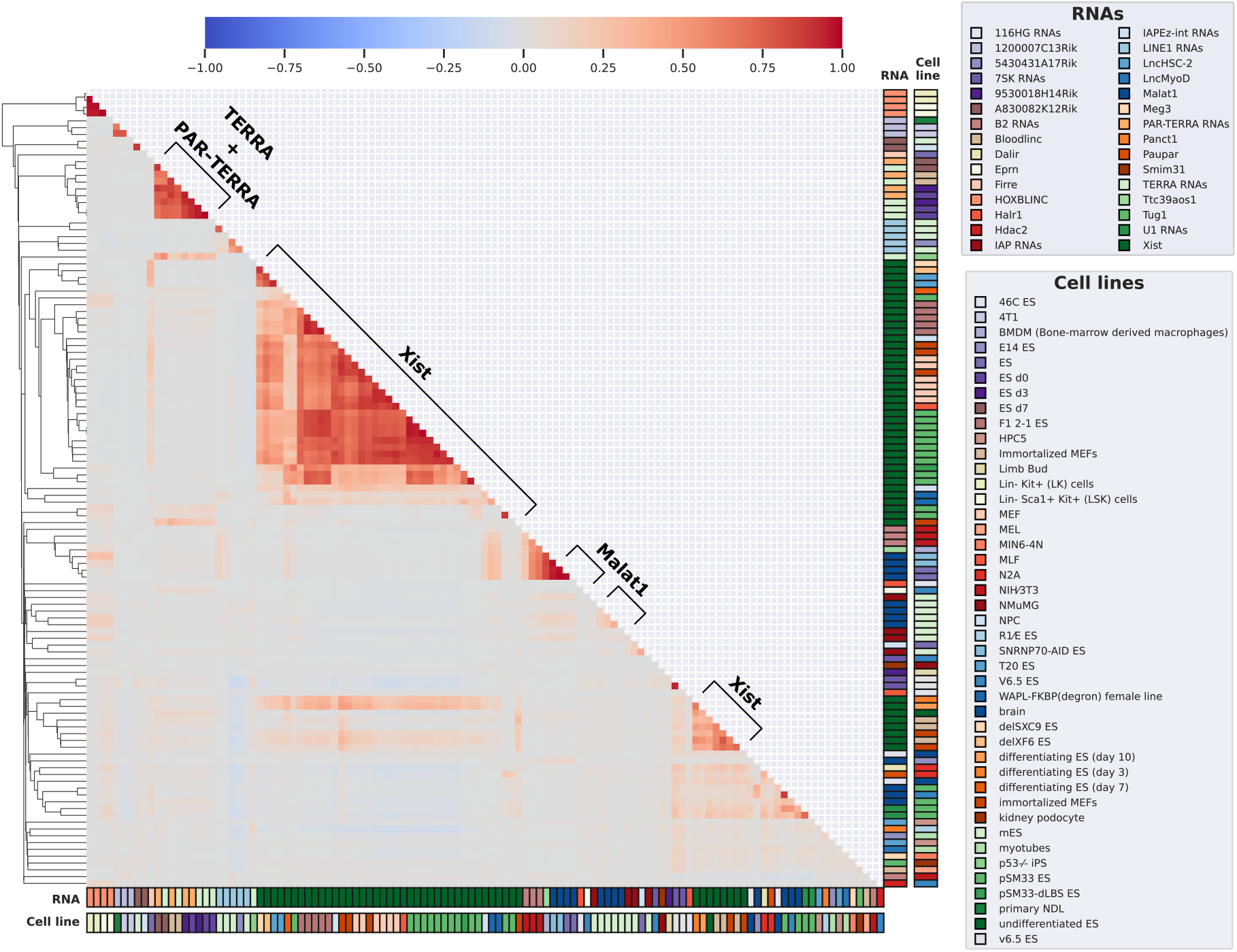
Pearson correlations between normalized contacts (that fall within the MACS2 peaks) tracks from OTA experiments on *Mus musculus* with merged replicates. Clustering by correlation. Genomic bin size: 10 kb, contacts filtering: >100 kb from the RNA source gene. Numerical values are given in Suppl. table 6.

Several large clusters for Xist RNAs as well as a cluster of TERRA and PAR-TERRA RNAs can be identified. The most conservative distribution of contacts on different cell lines is observed for Xist RNA. Its contacts tracks form a single cluster on raw and normalized data, and two separate clusters on peaks. The separation into two clusters in the latter case is explained by different additional treatments of the cells. For experiments from the smaller cluster, RNA Xist knockoff was performed followed by contacts recovery within a few hours. Also large clusters are formed by Malat1 RNA in *Mus musculus*, MALAT1 and NEAT1 in *Homo sapiens*. One of the Malat1 clusters also includes B2 and Ttc39aos1 RNAs. This can be explained by colocalization with CTCF binding sites observed for these three RNAs [21,22]. In addition, Meg3 RNA contacts tracks have a fairly good correlation with Xist tracks (Figure 5, pale yellow vertical line to the left of the Xist cluster). A weak negative correlation with Xist contacts is observed for the tracks of LINE1 RNAs contacts.

#### 3.3.2. Comparison of ATA experiments

Next, we assessed the correlations between total contact tracks from different ATA experiments for mouse and human (Figures 6, Suppl. Figure 4). After the normalization procedure, the correlation between experiments increases (Figures 6, Suppl. Figure 4: A, B), indicating the effectiveness of our normalization approach. At the same time, filtering contacts by distance from RNA source gene (>100 kb) did not lead to significant changes in correlations between experiments (Figures 6, Suppl. Figure 4: C). This is because although the density near RNA source genes is higher, the total number of distant contacts is much higher (Suppl. Table 3: column “Distant contacts (%)”). In contrast, the separation of all RNAs contacting chromatin into mRNAs and ncRNAs showed that mRNAs make the main contribution to the correlation between experiments (Figures 6, Suppl. Figure 4: E-F). This may be explained by the fraction of mRNA contacts in the mouse and human data being on the order of 70% except for experiments where additional cell treatments were applied (Suppl. Table 3, column “Share of mRNA contacts (%)”). There, NPM-experiments formed a separate cluster.

**Figure 6.**
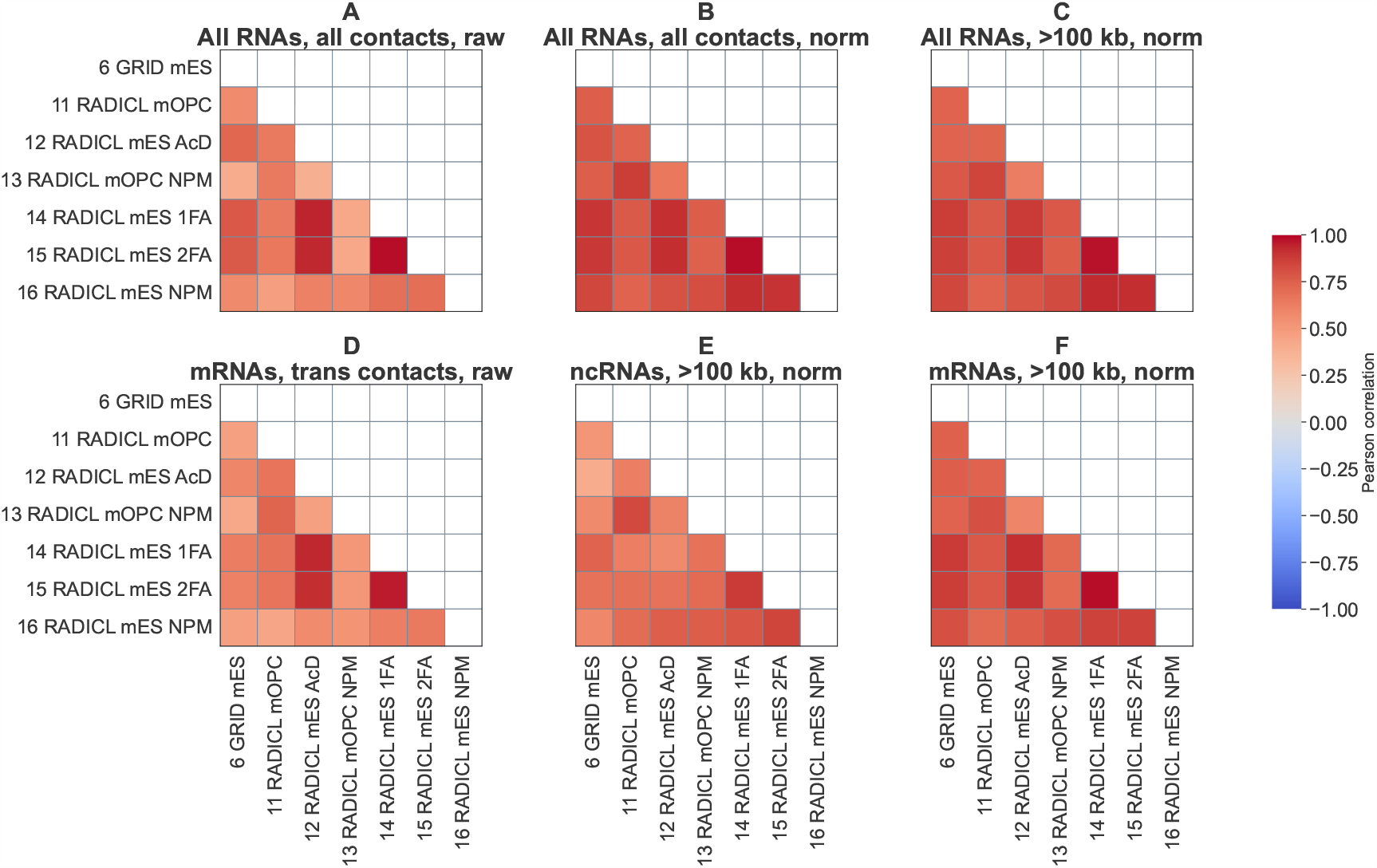
Pearson correlations between contacts tracks of ATA experiments, *Mus musculus*. (**A**) Raw contacts of all RNAs. (**B**) Background-normalized contacts of all RNAs. (**C**) Background-normalized contacts of all RNAs (contacts filtering: >100 kb from the RNA source gene). (**D**) Raw trans contacts of all mRNAs except for the 50 most contacting and 1000 least contacting mRNAs. (**E**) Background-normalized contacts of all ncRNAs (contacts filtering: >100 kb from the RNA source gene). (**F**) Background-normalized contacts of all mRNAs (contacts filtering: >100 kb from the RNA source gene). Genomic bin size: 10 kb. The format of the experiment labels is as in the figure scaling-ATA. The numerical values of correlations can be found in the Suppl. table 7.

The previous analysis considered the total number of contacts tracks. A more detailed analysis involves comparing the contacts tracks of individual RNAs from different experiments. To do this, we constructed similar heatmaps for some of the most contacting ncRNAs with chromatin (Figures 7 and Suppl. Figure 5). Mouse Malat1 and CT010467.1 RNA contact tracks correlate between all experiments. NPM-experiments (Exp.ID: 13, 16) and actinomycin D (Exp.ID: 12) are expectedly prominent for Kcnq1ot1 and Pvt1. Human MALAT1 and NEAT1 RNAs are also correlated in all experiments. The correlations of FTX and PVT1 contacts tracks from experiments with few contacts have low correlations with each other (Exp.IDs: 1, 2, 4, 5, 9). The correlations are affected by the type of treatment and number of contacts (Suppl. Figure 6) but are almost unaffected by cell type.

**Figure 7.**
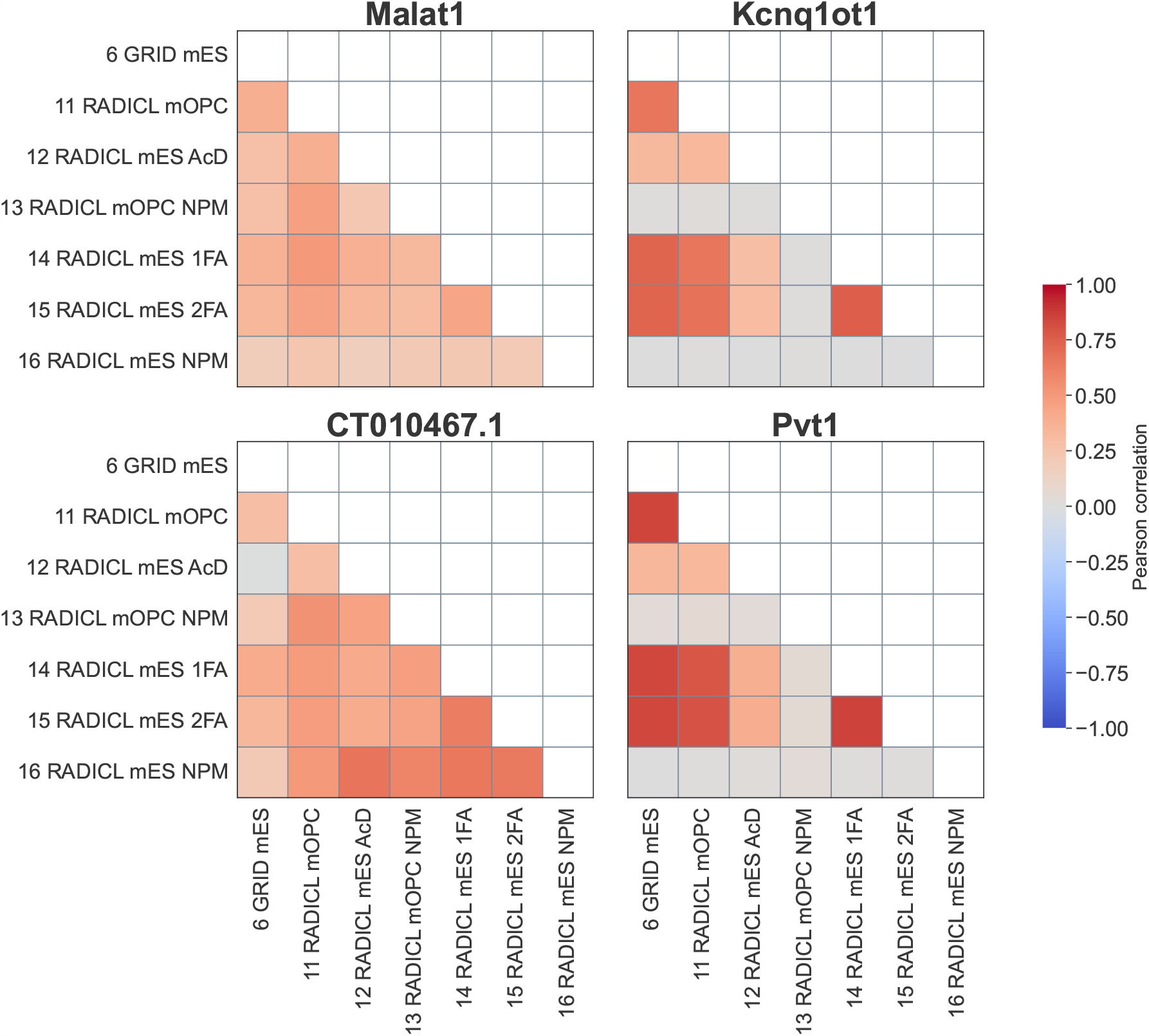
Pearson correlations between normalized contacts tracks of highly contacting ncRNAs from ATA experiments, *Mus musculus*. Genomic bin size = 10 kb, contacts filtering: >100 kb from the RNA source gene. The format of the experiment labels is as in the Figure 3. The numerical values of correlations and the normalized number of contacts in these tracks can be found in the Suppl. table 8.

#### 3.3.3. Comparison of ATA experiments with OTA experiments

ATA experiments allow us to infer the contacts of all RNAs in the cell at once, but the observed number of contacts for each individual RNA is much smaller than for OTA. This reduces both the completeness and accuracy of the results obtained. For each OTA experiment, the tracks of the corresponding RNAs were extracted from all ATA data.

The resulting tracks were normalized, and Pearson correlations were calculated for them (Figure 8). We used a bin size of 100 kb, as the density of contacts of each RNA is much lower in ATA data than in OTA data. With the use of OTA as a benchmark, the performance of ATA methods can be evaluated from the obtained correlations: Red-C and GRID were more accurate than RADICL and iMARGI. OTA experiments clustered by RNAs as expected. The correlations between OTA and ATA increase with the number of contacts of the considered RNA in ATA (Suppl. Figure 7). This means that the difference between the contacts counts for the considered RNAs from the OTA and ATA data may be due to the insufficient coverage of the RNAs in ATA. For example, the ncRNA MALAT1, which has a large number of contacts in all ATA experiments, shows a high correlation with OTA data (Figure 8). Other ncRNAs have significantly fewer contacts in ATA experiments, and their correlations with OTA are also lower. Surprisingly, although the GRID experiment shows only 237 contacts for ncRNA Xist, the correlation with most of the OTA data is still clearly visible. This seems to be due to the fact that although ATA data misses a large number of Xist RNA contacts, the observed ones are in good agreement with those from OTA, which cover chromosome X quite densely. The NPM-experiments (Exp.IDs: 13, 16) significantly lose correlation with OTA experiments. It is important to note again that OTA and ATA data have poor agreement across cell lines and additional chemical treatments (Suppl. Tables 1 and 2), which may also explain the lack of specific RNA contacts in ATA data.

**Figure 8.**
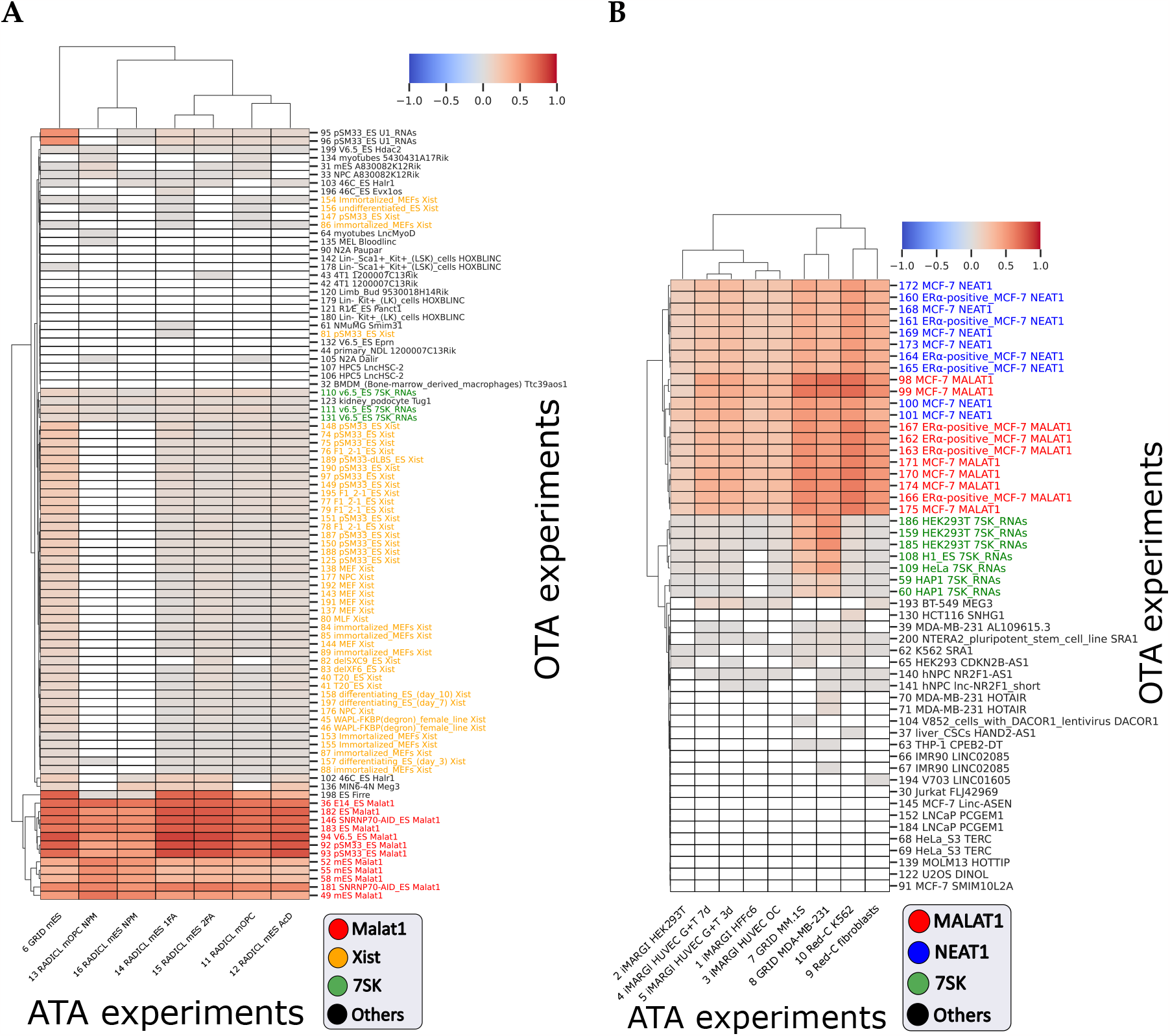
Pearson correlations between normalized contacts tracks from ATA and OTA experiments with merged replicates: (**A**) *Mus musculus*; (**B**) *Homo sapiens*. White color indicates correlations with *p − value >* 10^*−*2^ or the absence of contacts of the considered RNA in ATA data. Genomic bin size: 100 kb for OTA and ATA contacts tracks, contacts filtering: >100 kb from the RNA source gene. The format of the ATA experiment labels is as in the Figure 3. The format of the OTA experiment labels: *“Exp.ID from RNA-Chrom _ cell line _ RNA”*. In Suppl. tables 9 and 10 the numerical values are given.

### 3.4. Comparative analysis of the most contacting RNAs in ATA data

There are a number of RNAs that retain a significant number of interactions with chromatin in different cell lines due to their biological functions. For example, in each ATA experiment, the long non-coding RNA MALAT1 exhibits a high number of contacts. We tested whether the ranks of RNAs with highest frequencies of contacts would be preserved across different ATA experiments and whether this would depend on their biotype.

All RNAs from each ATA experiment were ordered by the normalized number of contacts (contacts filtering: >100 kb from the RNA source gene). Pairwise intersections of the corresponding top percentiles were taken to calculate the Jaccard measure defined as the ratio of the lists intersection size to the lists union size. There is a moderate (Jaccard index=0.3..0.6) overlap between the RNA tops in the pairs (Figure 9: A-B). However, for both human and mouse data, the sets of RNAs obtained as overlaps of all top 5% RNAs in terms of the number of contacts with chromatin (“RNA-leaders”) hold their high rank (a paired Wilcoxon test, Suppl. Figure 8: A-B). Statistical significance below the established level (*p− value <* 0.05) was observed in only two cases: Exp.ID 14 vs 16 and 15 vs 16, where the 16th experiment was performed by RADICL on mouse embryonic stem cells with protease K treatment (see Suppl. Figure 8B). This type of treatment dramatically alters the structure of contacts, therefore such experiments were excluded from further analysis, as well as the experiment with actinomycin D treatment. We also excluded iMARGI and “Red-C fibroblasts” experiments because of the small number of contacts in them. The new intersections of the top 5% RNAs in terms of the number of contacts with chromatin also retain their ranks in both human and mouse data (Suppl. Figure 8: C-D). Most of the RNA-leaders (1945 of 2036 in the case of human data and 1481 of 1540 in the case of mouse data) are protein-coding (Suppl. Figure 9: A-B), which will be discussed below. However, in addition to protein-coding RNAs, these intersections also contain various types of ncRNAs: long non-coding RNAs, small nuclear RNAs, small nucleus RNAs, antisense RNAs, and several other biotypes.

**Figure 9.**
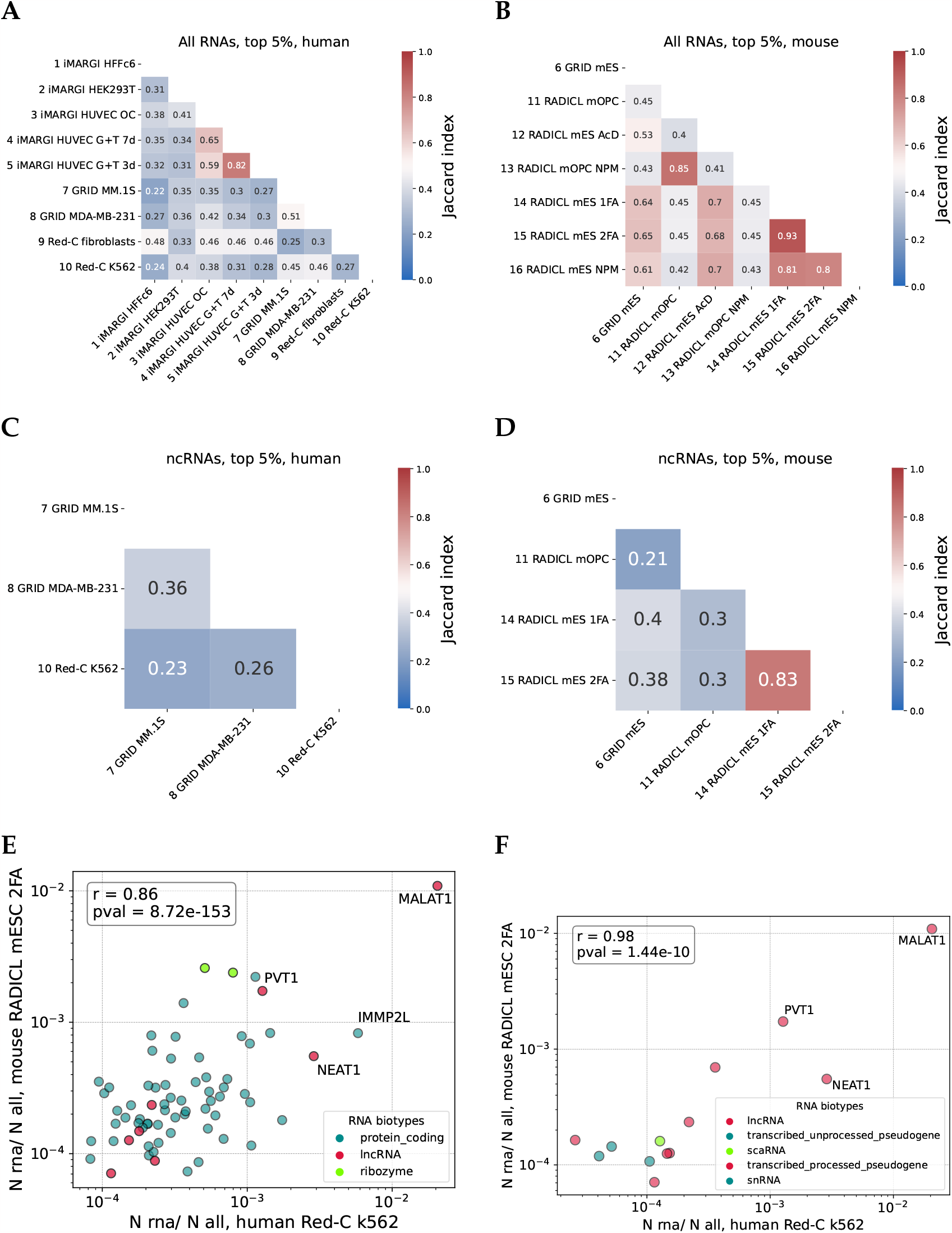
(**A, B**) Heat map of pairwise Jaccard measure values for RNA-leaders across all ATA experiments (**C, D**) and for ncRNA-leaders across comparable ATA experiments. The format of the ATA experiment labels is as in the figure scaling-ATA. (**E**) Comparison of contacts fractions for RNA-leaders orthologs with high ChP for human and mouse. (**F**) Comparison of contacts fractions for ncRNA-leaders orthologs with high ChP for human and mouse.

We compared the contactability (number of contacts) of RNA-leaders in human (“Red-C k562” experiment) with the contactability of their orthologs in mouse (“RADICL mESC 2FA” experiment). A fraction of human RNA-leaders have orthologs among mouse RNA-leaders (809 of 2036), including 2 ribozymes (RMRP_2 and RPPH1) and 8 long non-coding RNAs (MALAT1, NEAT1, PVT1, AL590627.3, DLEU2, LINC01004, TRAF3IP2-AS1, LAMTOR5-AS1), but the majority were protein-coding RNAs (799).

Chromatin potential characterizes the propensity of RNAs to contact chromatin. We selected RNA-leaders with high chromatin potential (ChP > 1.5) and their orthologs and compared the fraction of their contacts in “Red-C k562” and “RADICL mESC 2FA” data (Figure 9E). We observed high correlation (r=0.86, *p −value* = 8.72*e−* 153). Such RNAs retain a high degree of interaction with chromatin in various cell lines of different organisms.

In addition, we separately examined the top 5% overlap of ncRNAs only. In most cases, we observed a moderate Jaccard measure (*∼*0.3..0.6), except for the comparison of Exp.ID 14 vs 15, where the value is 0.83 (Figure 9: C, D). Interestingly, both of these experiments were performed on the same cell line, mouse embryonic stem cells. For the ncRNA-leaders, we found that the ranking was conserved (Suppl. Figure 8 E-F). The biotype diversity of ncRNA-leaders is shown in Suppl. Figure 10: A-B. The resulting correlation between contacts fraction of ncRNA-leaders in mouse and human was found to be strong (r=0.98, *p −value* = 1.44*e−* 10) (Figure 9F).

**Figure 10.**
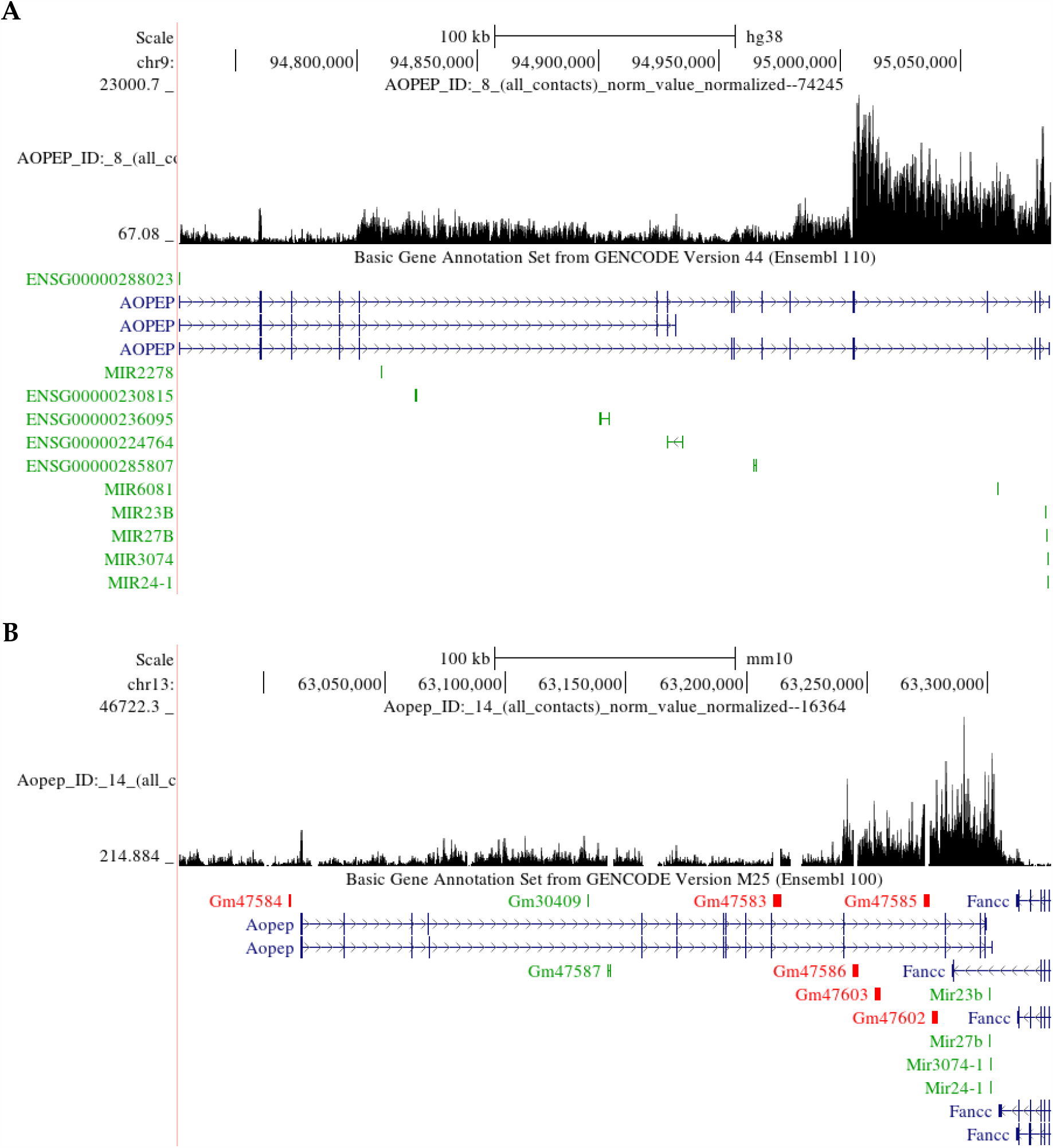
Distribution of (**A**) human and (**B**) mouse AOPEP RNA-parts of contacts across of the RNA source gene in Genome Browser [19]. Bin size: 200 nucleotides.

As noted previously, a significant proportion of the high-ranking positions in the various ATA experiments is occupied by protein-coding RNAs. However, the primary function of this RNA biotype does not appear to involve interactions with chromatin. In addition to high contactability, protein-coding RNA-leaders are also characterized by significantly elevated chromatin potential values (Suppl. Figure 9: C-D). It can be hypothesized that the annotated gene of a conserved highly contacting mRNA contains within it an unannotated gene of a ncRNA, or a fragment of this mRNA plays an independent regulatory role. In this case, we will observe an uneven distribution of RNA-parts of contacts across the RNA source gene, with the appearance of increased densities of RNA-parts mapping (hereafter referred to as “peaks”) in regions important for interaction with chromatin.

Let us consider only those mRNA-leaders in which both orthologs have a chromatin potential exceeding 1.5. The number of such mRNAs is 59. Uneven distribution of RNA-parts of contacts is observed in most of the selected mRNAs. For example, the mRNA AOPEP has a pronounced peak at the 3’-end of both the human gene and its mouse ortholog (Figure 10). It should be noted that the peaks are not additionally annotated. All of the above factors point to the possibility that these peaks contain ncRNAs that have not been previously annotated or that there is a regulatory role for the corresponding mRNA site. However, more detailed analysis is needed to confirm this assumption.

## 4. Discussion

The RNA-Chrom database collected all RNA-chromatin interactions data for human and mouse available by early 2022: 171 OTA experiments (118 for mouse and 53 for human) and 16 ATA experiments (7 for mouse and 9 for human). The aim of the present work was primarily to assess the completeness and quality of the interactome data. For this purpose, we performed a data comparative analysis.

Firstly, the replicates in the experiments had to be consistent. The comparison of replicates already showed that the number of reads is a fundamentally important parameter, especially for ATA experiments. Moreover, for individual RNAs, the consistency of replicates is maintained only for RNAs with a sufficiently large number of contacts. Hence, an important conclusion is that the data can be considered sufficiently complete only for those RNAs that have a large number of contacts. To increase the signal, replicates can be merged, but this approach may significantly increase not only the signal but also the noise. In the ChIRP-seq [4] method (OTA experiments), authors used an approach based on the intersection of similar data obtained by “odd” and “even” probes. This approach should significantly reduce noise. However, a significant portion of the signal will be lost, especially if at least one of the replicates has low coverage, so this approach is not readily applicable to the ATA data.

The analyzed data have limited precision by their nature, so data binning is often used when analyzing RNA-chromatin interactome data. The choice of bin size can significantly affect the results. Comparison of replicates allowed us to estimate the appropriate genomic bin size (Suppl. Figure 11). It appears that the optimal bin size depends on the sequencing depth. For the data we considered in this paper, suitable bins are 10 kb for OTA data and 100 kb for ATA data. However, for some types of analysis, variable-sized bins can be used. Previously, it was noted that the RNA cis-contacts density decreases with the distance from an RNA source gene (scaling). Our analysis showed that this dependence is present in almost all datasets, both ATA and OTA, barring NPM-experiments. The nature of this dependence differs somewhat between experiments and for different RNAs. It reflects not only the polymerase trace but also some features of chromatin folding, as well as RNA diffusion rates from the site of transcription. This bias also needs to be taken into account when analyzing the data, which is why we did not consider the RNA contacts closer than 100 kb to the RNA source gene.

Another source of bias is chromatin accessibility. Less accessible regions of chromatin are less susceptible to cleavage by restrictases, DNAases, and ultrasound, so generally fewer contacts are observed in heterochromatin regions. To account for this bias, normalization to background was applied. Finally, non-specific interactions result in additional noise, which may be reduced by peak calling. We compared the data from OTA experiments. Our analysis showed that the control for the mentioned factors is necessary. Peak calling gives the largest contribution, and polymerase trace suppression gives the smallest contribution. As a result of comparing OTA data, we obtained a quite expected result – the data are clustered well by RNA biotypes. Moreover, several clusters corresponding to different experimental conditions and cell lines are distinguished for Xist. Comparison of ATA experiments with each other showed, first of all, that the correlation of the data is affected by sequencing depth and types of additional cell treatment in the experiments. No significant differences were found between cell lines neither in ATA data nor in OTA data. Comparison of ATA data with OTA data showed that the correlation of their contacts maps was also primarily influenced by cell processing and the number of contacts.

We compared the lists of RNAs with the highest densities of contacts with chromatin in ATA data. It turned out that for most pairs of experiments the lists of the most contacting RNAs are similar. At the same time, the majority of RNAs in the top 5% in terms of the number of contacts with chromatin are mRNAs. This happens primarily due to the fact that mRNA has a much higher expression.

Our interspecies comparison revealed that a significant proportion of highly contacting RNAs in human are also highly contacting in mouse. This is true for both mRNAs and ncRNAs. Comparison of the distributions of mRNA-parts of contacts across RNA source genes showed that there are highly contacting regions on mRNAs, and these distributions are quite similar in human and mouse orthologs. It can be assumed that these regions correspond to previously unannotated non-coding RNAs.

To our knowledge, only a single study [20] attempted to combine all available OTA data by the year 2019 to analyze together and with a large amount of epigenetic data. Zhang et al. analyzed 23 OTA datasets, 12 for human and 10 for mouse, but they did not consider ATA data, the number of which was only 10 datasets at that time. In our work, we performed a comparative analysis of a much larger number of data and also examined ATA data. Our correlations between RADICL and GRID ATA experiments correspond to the results obtained by the authors of the RADICL-seq method [12]. We aimed to assess data quality and completeness to establish a set of good quality-control practices for further qualitative studies.

Our results may help researchers not only to perform higher quality experiments, but also to use RNA-chromatin interactome data more effectively in integrative analysis with other genome-wide data, such as chromatin structure, gene expression, localization of DNA-binding and chromatin-modifying proteins, or epigenetic data. We believe that RNA-chromatin interactome data, despite being noisy, are crucial for understanding functions of non-coding RNAs.

## Supplementary Materials

The following supporting information can be downloaded at: https://www.mdpi.com/article/10.3390/biom1010000/s1, Figure S1: Pearson correlation between human replicates as a function of the minimum number of raw contacts in them; Figure S2: Pearson correlations between contacts tracks from OTA experiments with merged replicates; Figure S3: Pearson correlations between normalized contacts (that fall within the MACS2 peaks) tracks from OTA experiments on *Homo sapiens* with merged replicates; Figure S4: Pearson correlations between contacts tracks of ATA experiments, *Homo sapiens*; Figure S5: Pearson correlations between normalized contacts tracks of highly contacting ncRNAs from ATA experiments, *Homo sapiens*; Figure S6: Number of contacts in pairs of ATA experiments: (**A**) mouse and (**B**) human; Figure S7: Pearson correlation (*p − value >* 10^*−*2^) between normalized contacts tracks of RNAs from OTA experiments and contacts tracks of the corresponding RNAs from ATA experiments as a function of the number of contacts in the corresponding ATA tracks; Figure S8: Heat map of paired Wilcoxon test p-values; Figure S9: Diversity of RNA-leader biotypes in comparable ATA experiments and distribution of chromatin potential values; Figure S10: Diversity of ncRNA biotypes from the intersection of top 5% ncRNAs only in terms of the number of contacts with chromatin from comparable ATA experiments; Figure S11: Average Pearson correlation across replicates of raw ATA data depending on genomic bin size; Table S1: Diversity of OTA and ATA (Homo sapiens) data in the RNA-Chrom database; Table S2: Diversity of OTA and ATA (Mus musculus) data in the RNA-Chrom database; Table S3: ATA data description; Table S4: OTA data description; Table S5: Representation of different RNA biotypes in the “top 25% of cis-contacting RNAs” sample for the corresponding ATA experiment; Table S6: Pearson correlations between OTA experiments on Mus musculus; Table S7: Pearson correlation between contacts tracks in ATA experiments on Mus musculus; Table S8: Pearson correlation between contacts tracks of RNAs (Malat1, Kcnq1ot1, CT010467.1, Pvt1) in ATA experiments on Mus musculus; Table S9: Pearson correlation between OTA experiments and their corresponding RNA tracks from ATA experiments on Mus musculus; Table S10: Pearson correlation between OTA experiments and their corresponding RNA tracks from ATA experiments on Homo sapiens; Table S11: Pearson correlations between OTA experiments on Homo sapiens; Table S12: Pearson correlation between contacts tracks in ATA experiments on Homo sapiens; Table S13: Pearson correlation between contacts tracks of RNAs (MALAT1, NEAT1, FTX, PVT1) in ATA experiments on Homo sapiens.

## Author Contributions

GR, scaling analysis, manuscript preparation; AV, correlation analysis of replicates, correlation analysis of OTA data and comparison of ATA data with OTA data; VS, correlation analysis of replicates for individual RNAs and comparison of ATA data; LG, leader RNA analysis and interspecies comparison; AM, general supervision, manuscript preparation.

## Acknowledgments

We thank Mikhail Moldovan and Herman Ashniev, who carefully read the manuscript and made helpful suggestions, and Kate Ivanova for English proofreading.

## Disclaimer/Publisher’s Note

The statements, opinions and data contained in all publications are solely those of the individual author(s) and contributor(s) and not of MDPI and/or the editor(s). MDPI and/or the editor(s) disclaim responsibility for any injury to people or property resulting from any ideas, methods, instructions or products referred to in the content.

