## Supplementary Materials: for "Comparative analysis of the RNA-chromatin interactions data. Completeness and accuracy"

### 1. Supplementary figures

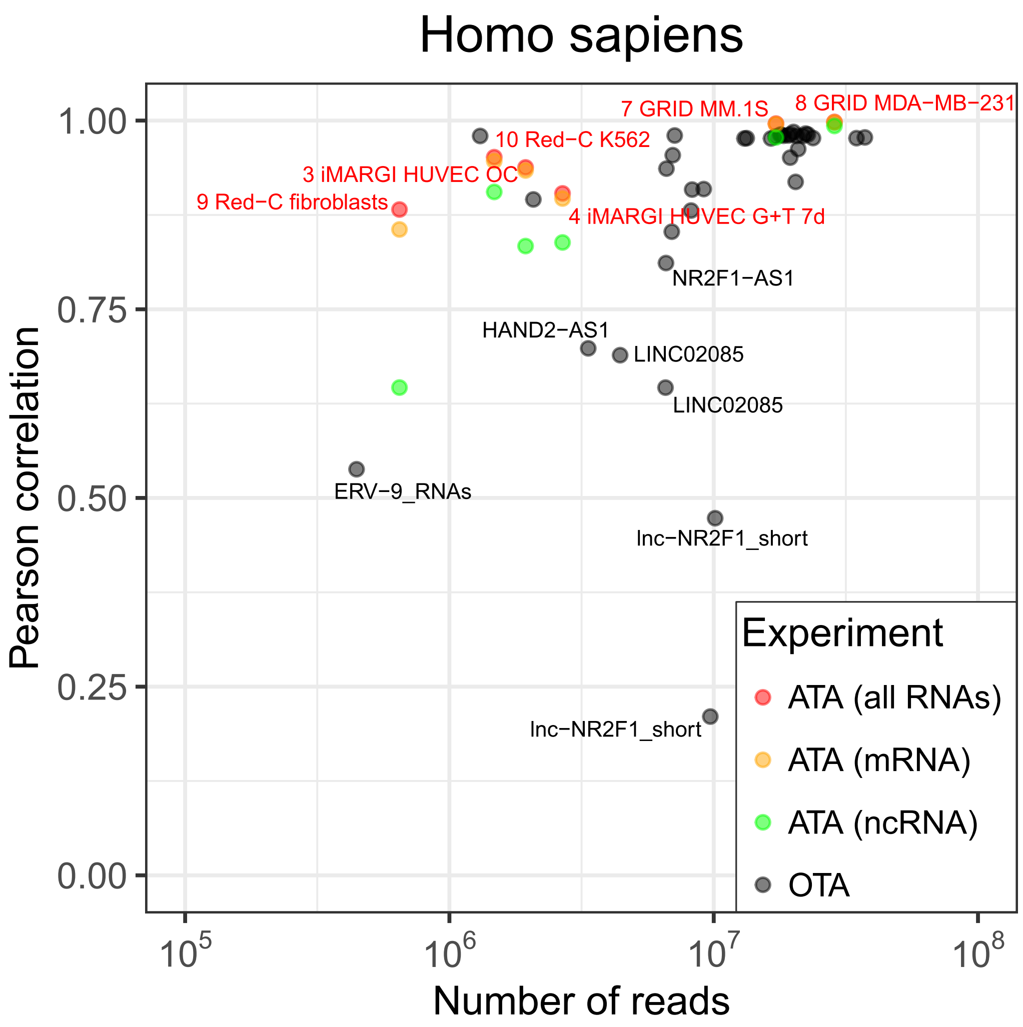


**Figure S1.** Pearson correlation between human replicates as a function of the minimum number of raw contacts in them. If an experiment had more than two replicates, the average of all pairwise correlations was taken. Genomic bin size: 10 kb (OTA experiments), 100 kb (ATA experiments), contacts filtering: >100 kb from the RNA source gene.

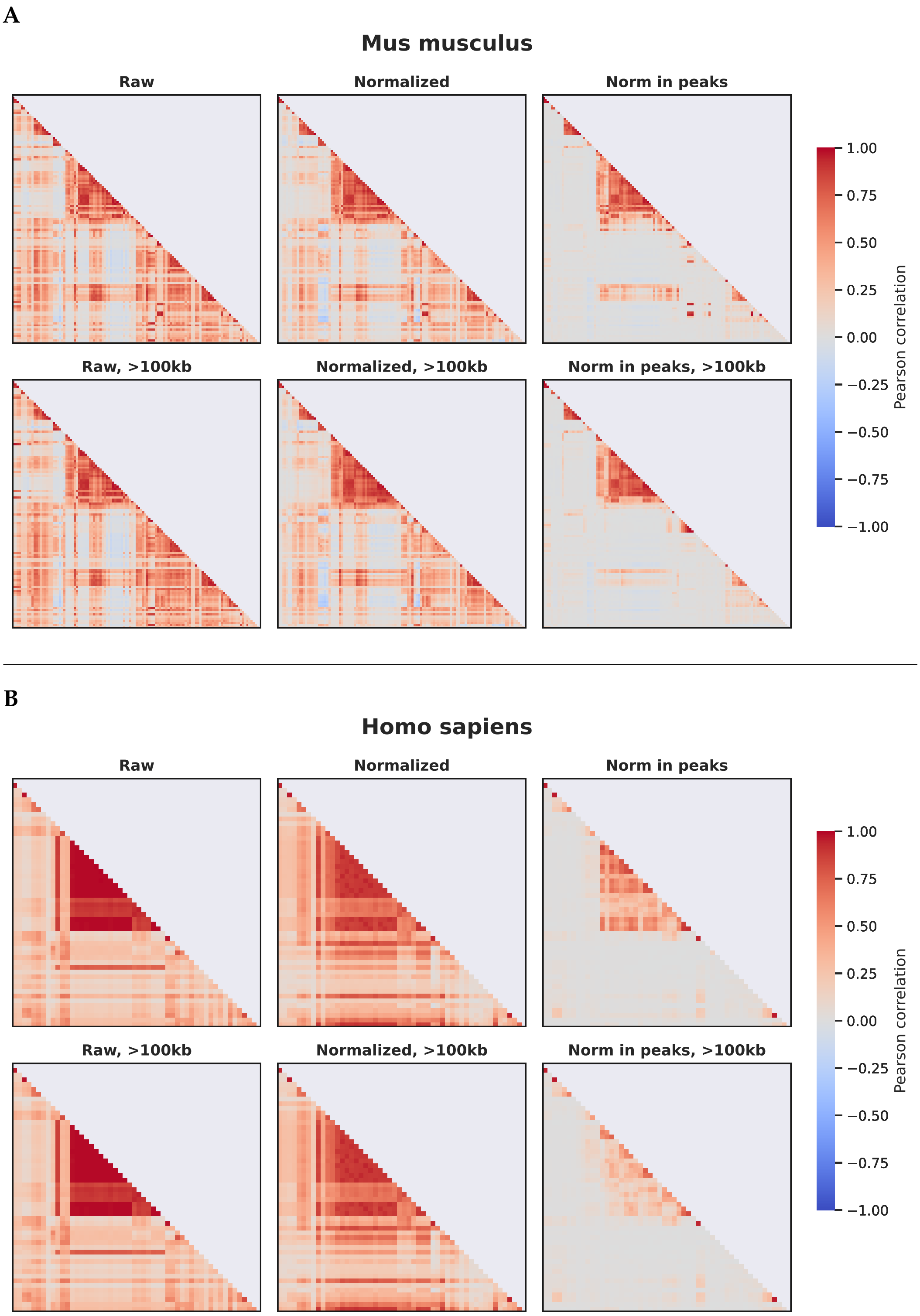


**Figure S2.** Pearson correlations between contacts tracks from OTA experiments with merged replicates. Genomic bin size: 10 kb. (**A**) For each processing stage of mouse OTA data with and without contacts filtering: >100 kb from the RNA source gene. The order of experiments is the same as in Figure 5 of the main text. (**B**) For each processing stage of human OTA data with and without contacts filtering: >100 kb from the RNA source gene. The order of experiments is the same as in Figure S3.

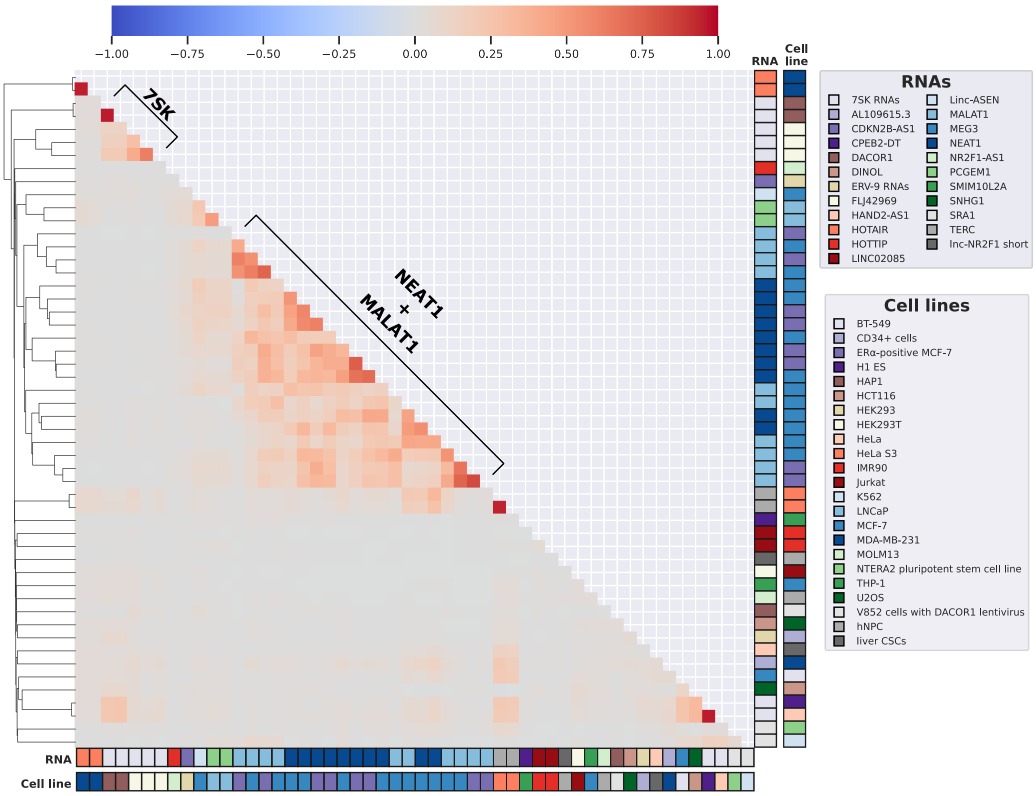


**Figure S3.** Pearson correlations between normalized contacts (that fall within the MACS2 peaks) tracks from OTA experiments on *Homo sapiens* with merged replicates. Clustering by correlation. Genomic bin size: 10 kb. Numerical values are given in Suppl. table 11.

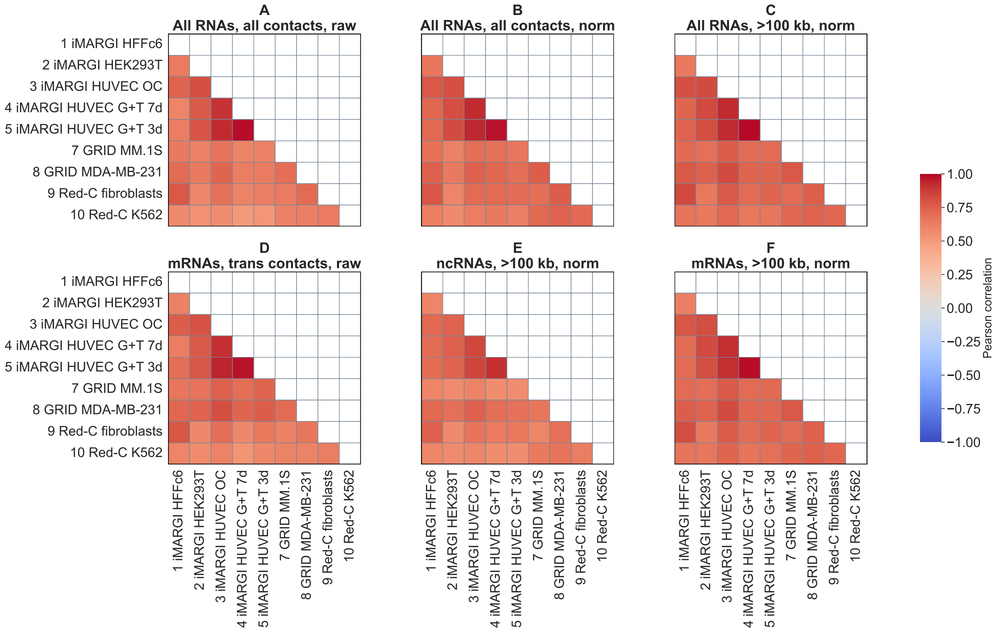


**Figure S4.** Pearson correlations between contacts tracks of ATA experiments, *Homo sapiens*. (**A**) Raw contacts of all RNAs. (**B**) Background-normalized contacts of all RNAs. (**C**) Background-normalized contacts of all RNAs (contacts filtering: >100 kb from the RNA source gene). (**D**) Raw trans contacts of all mRNAs except for the 50 most contacting and 1000 least contacting mRNAs. (**E**) Background-normalized contacts of all ncRNAs (contacts filtering: >100 kb from the RNA source gene). (**F**) Background-normalized contacts of all mRNAs (contacts filtering: >100 kb from the RNA source gene). Genomic bin size: 100 kb. The format of the experiment labels is as in the Figure 3 of the main text. The numerical values of correlations can be found in the Suppl. table 12.

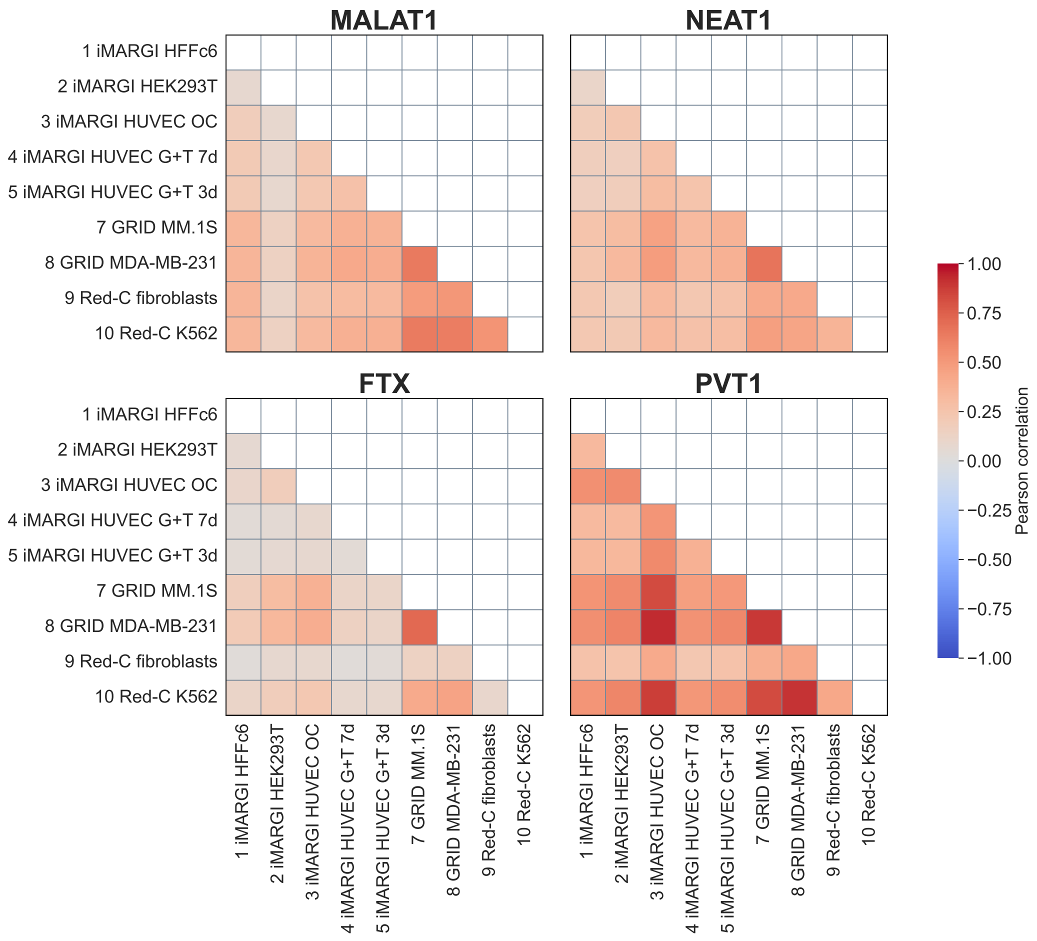


**Figure S5.** Pearson correlations between normalized contacts tracks of highly contacting ncRNAs from ATA experiments, *Homo sapiens*. Genomic bin size = 100 kb, contacts filtering: >100 kb from the RNA source gene. The format of the experiment labels is as in the Figure 3 of the main text. The numerical values of correlations and the normalized number of contacts in these tracks can be found in the Suppl. table 13.

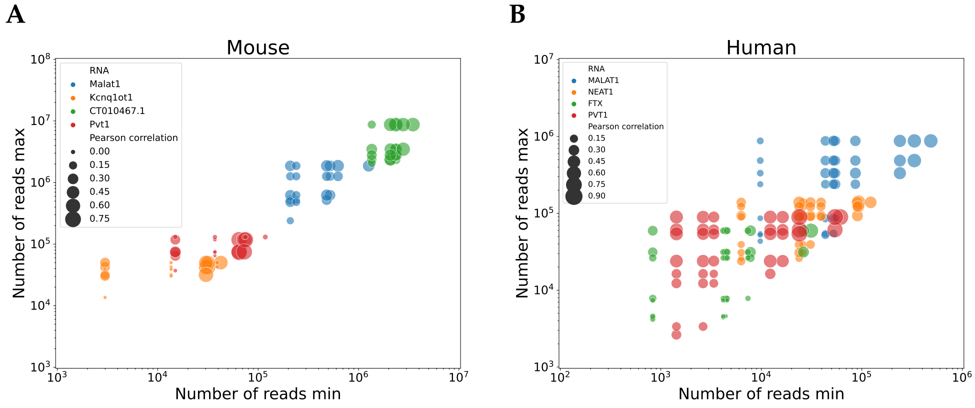


**Figure S6.** Number of contacts in pairs of ATA experiments: (**A**) mouse and (**B**) human. The vertical axis shows the number of contacts in the experiment with the maximum number of contacts, while the horizontal axis shows the number of contacts in the experiment with the minimum number of contacts. Each dot is the selected RNA in a pair of experiments. Color - RNA name, circle size - correlation between contacts tracks.

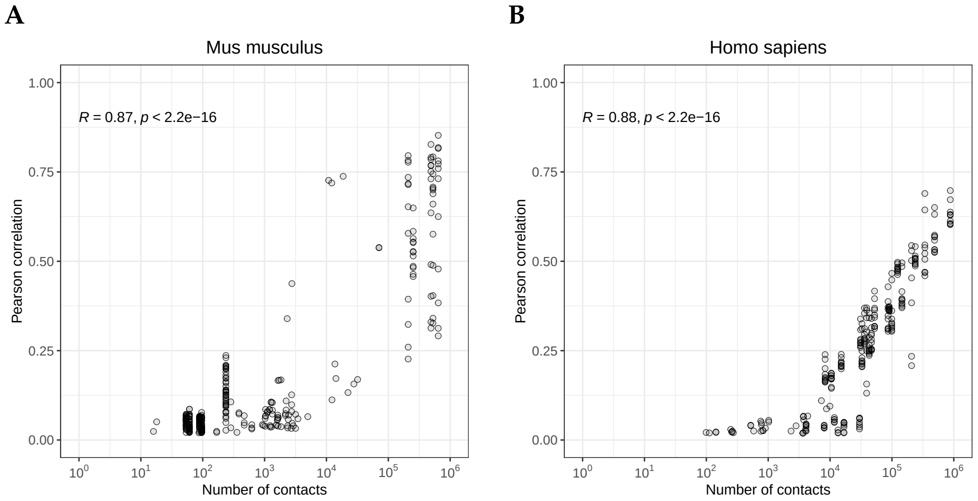


**Figure S7.** Pearson correlation ($p - value > 10^{-2}$) between normalized contacts tracks of RNAs from OTA experiments and contacts tracks of the corresponding RNAs from ATA experiments as a function of the number of contacts in the corresponding ATA tracks. (**A**) *Mus musculus*. (**B**) *Homo sapiens*.

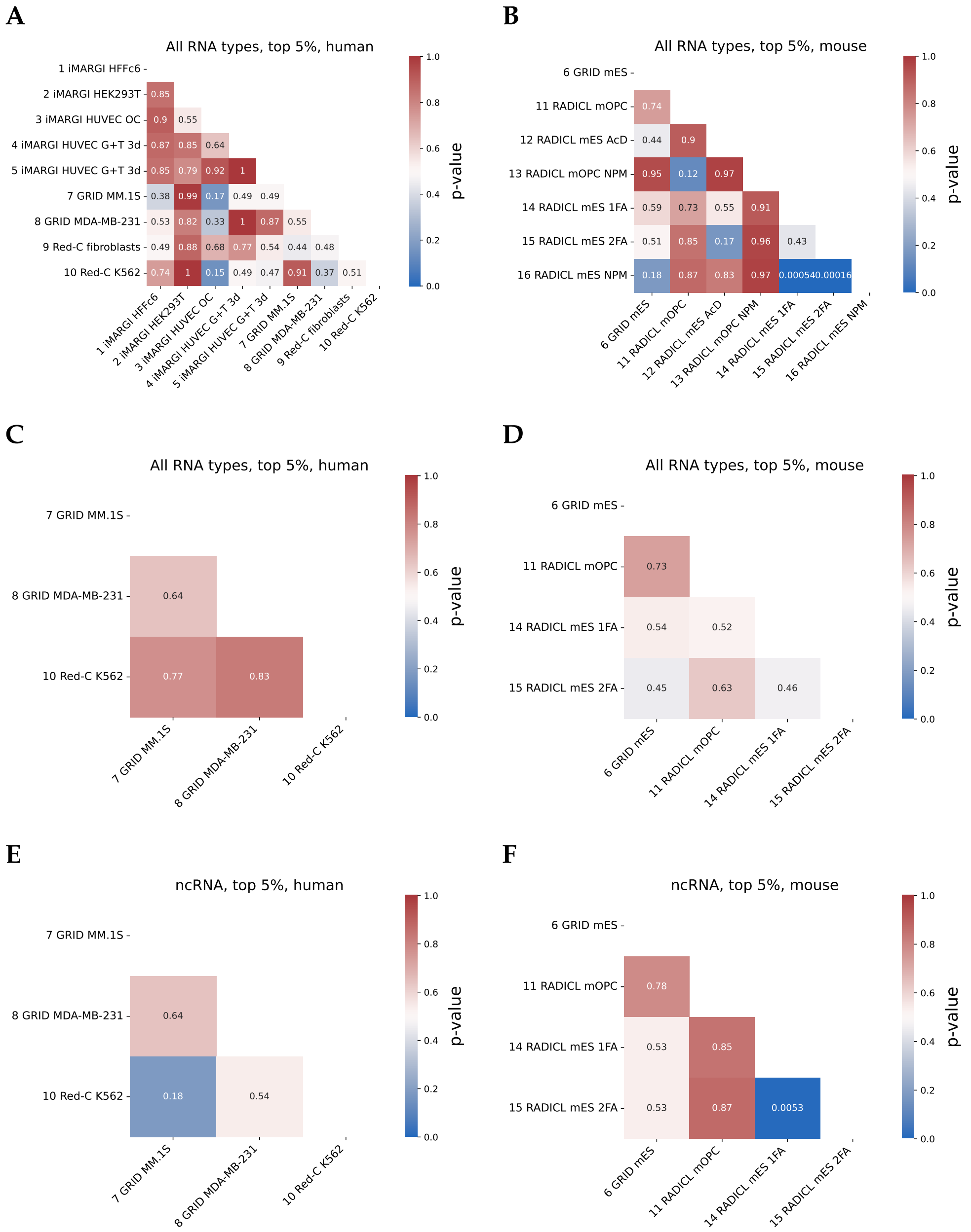


**Figure S8.** Heat map of paired Wilcoxon test p-values. (**A, B**) For RNAs from the intersection of tops 5% of all human and mouse ATA data. (**C, D**) For RNAs from the intersection of tops 5% of the comparable human and mouse ATA data. (**E, F**) For ncRNAs from the intersection of tops 5% of the comparable human and mouse ATA data. The format of the ATA experiment labels is as in the Figure 3 of the main text.

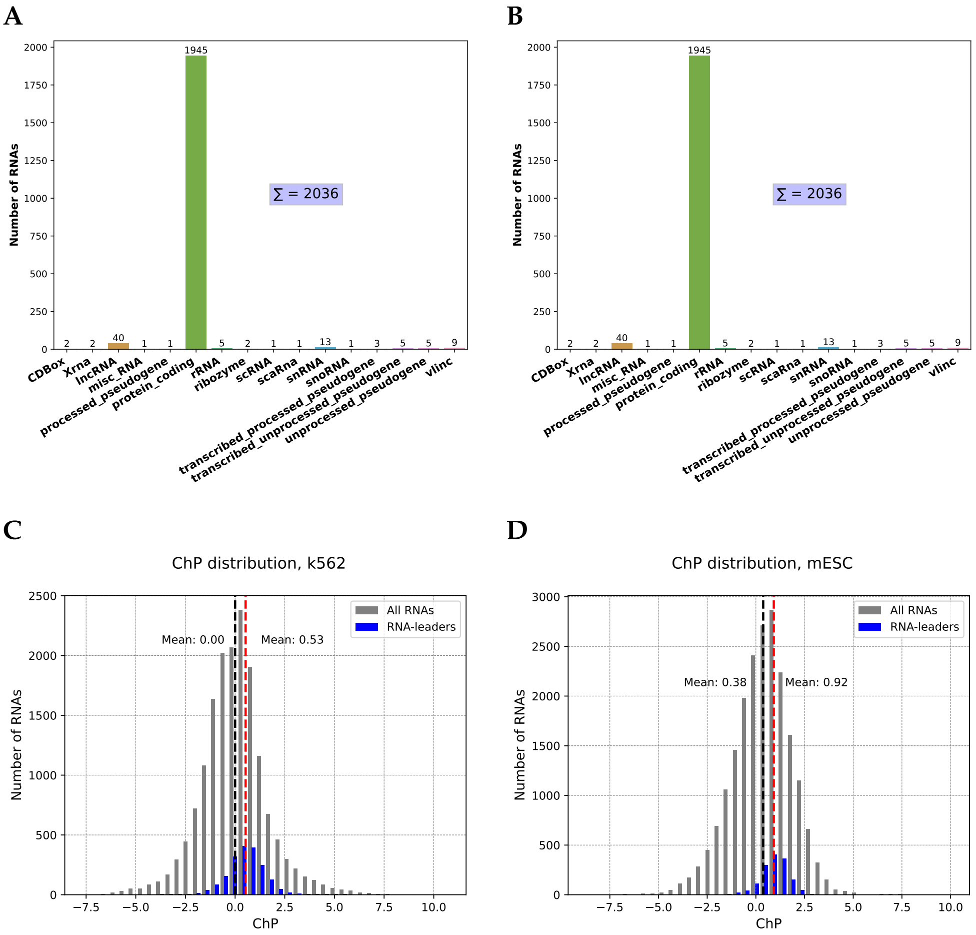


**Figure S9.** (**A, B**) Diversity of RNA-leader biotypes in comparable ATA experiments. (**C, D**) Distribution of chromatin potential values (ChP) of protein-coding RNA-leader from "Red-C k562" and "GRID mESC" experiments. Gray color – distribution of ChP of all protein-coding RNAs from the experiment. Blue color – the same distribution, but only of the highly contacting protein-coding RNAs.

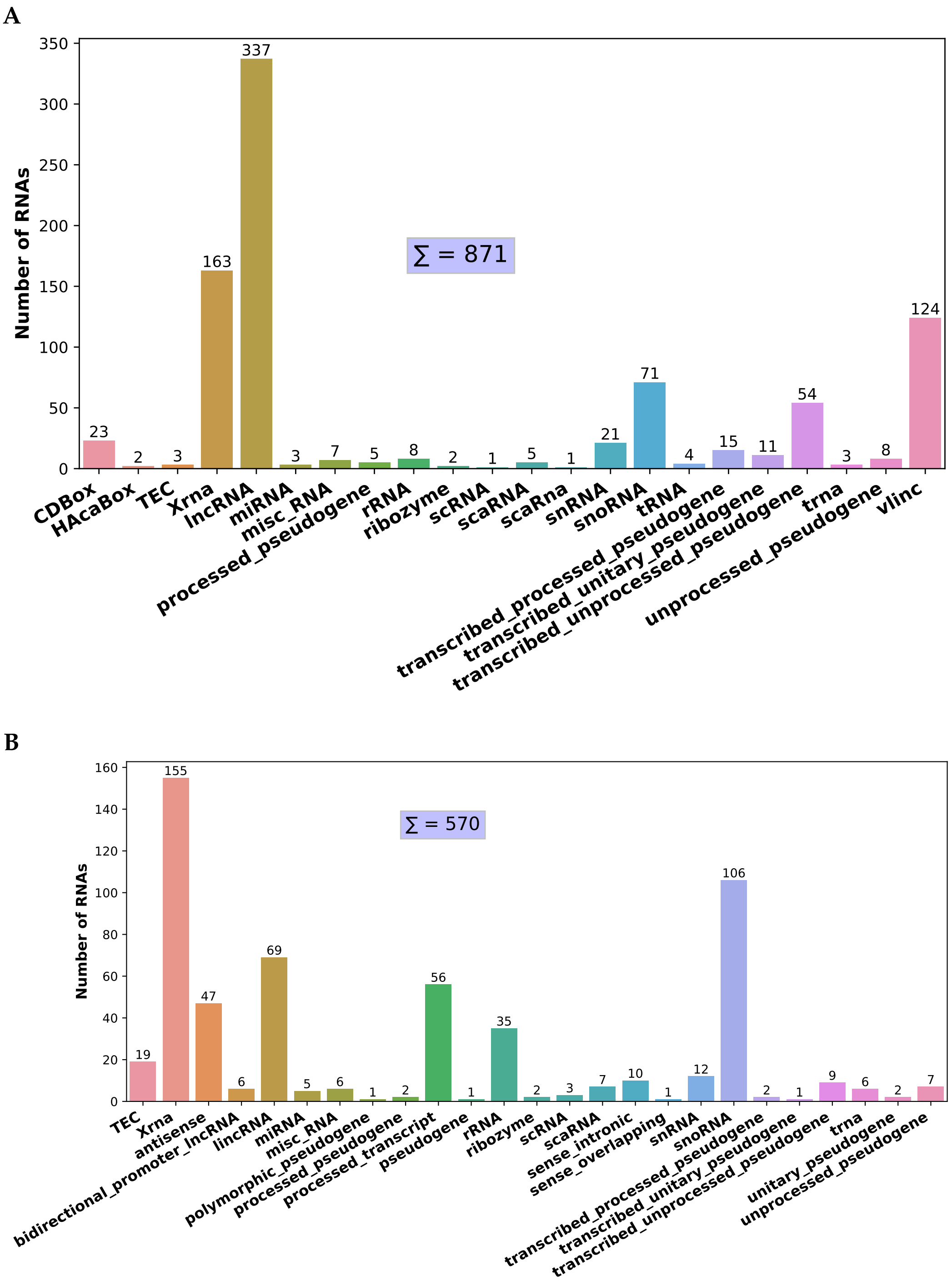


**Figure S10.** Diversity of ncRNA biotypes from the intersection of top 5% ncRNAs only in terms of the number of contacts with chromatin from comparable ATA experiments. (**A**) *Homo sapiens*; (**B**) *Mus musculus*.

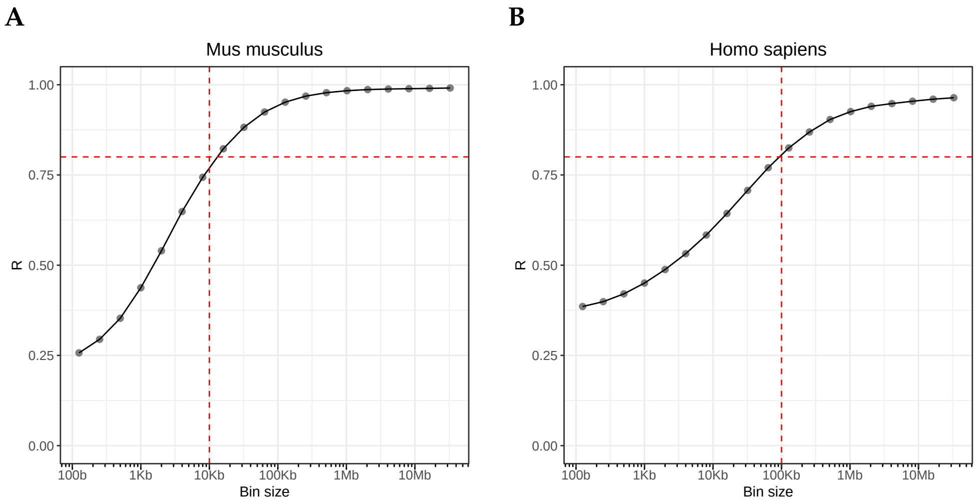


**Figure S11.** Average Pearson correlation across replicates of raw ATA data depending on genomic bin size. Contacts filtering: >100 kb from the RNA source gene. (**A**) *Mus musculus*; (**B**) *Homo sapiens*. The horizontal line indicates a correlation of 0.8, the vertical line indicates the corresponding bin size.

### 2. Supplementary Tables

1. **Supplementary Table 1.** Diversity of OTA and ATA (*Homo sapiens*) data in the RNA-Chrom database. N* – amount of experiments in the RNA-Chrom DB. Cell lines, for which both OTA and ATA data are available, are highlighted in green.
2. **Supplementary Table 2.** Diversity of OTA and ATA (*Mus musculus*) data in the RNA-Chrom database. N* – amount of experiments in the RNA-Chrom DB. Cell lines, for which both OTA and ATA data are available, are highlighted in green.
3. **Supplementary Table 3.** ATA data description.
4. **Supplementary Table 4.** OTA data description.
5. **Supplementary Table 5.** Representation of different RNA biotypes in the "top 25% of cis-contacting RNAs" sample for the corresponding ATA experiment.
6. **Supplementary Table 6.** Pearson correlations between OTA experiments on *Mus musculus*. Correlations are shown in the lower triangle and corresponding p-values in the upper.
7. **Supplementary Table 7.** Pearson correlation between contacts tracks in ATA experiments on *Mus musculus*.
8. **Supplementary Table 8.** Pearson correlation between contacts tracks of RNAs (Malat1, Kcnq1ot1, CT010467.1, Pvt1) in ATA experiments on *Mus musculus*. And also the normalized number of contacts in these tracks, contacts filter: >100 kb from the RNA source gene. Genomic bin size: 10 kb.
9. **Supplementary Table 9.** Pearson correlation between OTA experiments and their corresponding RNA tracks from ATA experiments on *Mus musculus*.
10. **Supplementary Table 10.** Pearson correlation between OTA experiments and their corresponding RNA tracks from ATA experiments on *Homo sapiens*.
11. **Supplementary Table 11.** Pearson correlations between OTA experiments on *Homo sapiens*. Correlations are shown in the lower triangle and corresponding p-values in the upper.
12. **Supplementary Table 12.** Pearson correlation between contacts tracks in ATA experiments on *Homo sapiens*.
13. **Supplementary Table 13.** Pearson correlation between contacts tracks of RNAs (MALAT1, NEAT1, FTX, PVT1) in ATA experiments on *Homo sapiens*. And also the normalized number of contacts in these tracks, contacts filter: >100 kb from the RNA source gene. Genomic bin size: 10 kb.

**Disclaimer/Publisher's Note:** The statements, opinions and data contained in all publications are solely those of the individual author(s) and contributor(s) and not of MDPI and/or the editor(s). MDPI and/or the editor(s) disclaim responsibility for any injury to people or property resulting from any ideas, methods, instructions or products referred to in the content.
